# Antibacterial effects of nanoimprinted moth-eye film in practical settings

**DOI:** 10.1101/325837

**Authors:** Miho Yamada, Kiyoshi Minoura, Takashi Mizoguchi, Kenichiro Nakamatsu, Tokio Taguchi, Takuya Kameda, Miho Sekiguchi, Tatsuo Suzutani, Shinichi Konno

## Abstract

Recent studies report that surfaces displaying micrometer-or nanometer-sized undulating structures exhibit antibacterial effects. In previous work, we described the use of an advanced nanofabrication technique to generate an artificial biomimetic moth-eye film by coating a polyethylene terephthalate (PET) film with nanoscale moth-eye protrusions made from a hydrophilic resin. This moth-eye film exhibited enhanced antibacterial effects in *in vitro* experiments. The aim of the present study was to verify the antibacterial efficacy of the Moth-eye film in practical environments. Three types of films (Moth-eye film, Flat film, and PET film) were used to compare antibacterial effects. Sample films were pasted onto hand-wash sinks at the testing locations. After several hours of elapsed time, bacteria from the surface of sample films were collected using one of three kinds of culture media stampers (to permit identification of bacterial species). The stampers were incubated for 48 hours at 35 °C, and the numbers of colonies were counted.

The number of common bacteria including *E. coli* and *S. aureus* from the Moth-eye film was significantly lower than that from the PET film (p<0.05) and that from the Flat film at 1 hour (p<0.05). This study found that the Moth-eye film had durability of antibacterial effect and the Moth-eye structure (PET coated with nanoscale cone-shaped pillars) had a physical antibacterial effect from the earlier time points. Therefore, the Moth-eye film might be useful for general-purpose applications in practical environments.

## INTRODUCTION

A large number of patients, ranging from 648 thousand to 1700 thousand patients per year in the United States, suffer from healthcare-associated infections (HCAIs) [1]. Continuing efforts are being made to better understand and control the spread of various infectious diseases. The prevention of hospital infections starts with hand-washing. Frequent sterilization and/or disinfection of environmental surfaces are also highly important. However, it is impossible to maintain hygienic cleanliness over the entire area of medical institutions where bacterial carriers and the readily infected are expected to mix. The situation has become more serious than ever with the appearance of drug-resistant bacteria such as methicillin-resistant *Staphylococcus aureus* (MRSA), vancomycin-resistant *Enterococcus* (VRE), and multi-drug-resistant *Pseudomonas aeruginosa* (MDRP). Antimicrobial technologies have been developed in response to strong demands for steady and operable actions in the field against hospital infections. In addition, one of the next development targets is incorporation of antibacterial materials into advanced medical equipment, where such materials would need to exhibit high stability and safety.

Various mechanisms for infection control have been widely used, i.e. chemical agents, exposure to high temperature, high pressure and ultraviolet rays, and incorporation of photocatalysts and so on. Silver particles [2] are among the more useful sources of antibacterial action. Materials that include silver particles are regarded as safe and durable, and can help prevent the growth of drug-resistant bacteria. Therefore, this material has been incorporated into a wide variety of environments, ranging from household products to medical equipment.

Recent studies have reported that surfaces displaying micrometer-or nanometer-sized undulations exhibit physical antibacterial effects. For instance, the wings of dragonflies and cicadas have been reported to have the ability to kill bacteria [3–5]. Shark skin [6] and sacred lotus leaf [7] have the ability to prevent the attachment of bacteria. These structures exhibit regular or random undulations ranging in size from several tens of nanometers to several micrometers. Artificial nanostructured surfaces having a regular array of pillars, approximately 200-nm high and spaced approximately 170 nm apart, also have been reported to have physical bactericidal effects [8].

We have developed an artificial moth-eye film that is fabricated using an advanced nano-imprinting technique. The resulting film possesses unique nanostructure arrays that exhibit a variety of useful functions, including super-hydrophilic or super-hydrophobic properties due to the larger surface area of the moth-eye film compared to a flat surface. In previous work, we reported that our moth-eye film exhibited enhanced antibacterial effects in *in vitro* experiments [9]. However, the moth-eye film has not been evaluated for antibacterial effects in practical environments. Hand-wash sinks are considered to be a major source of hospital infection [10], due to environmental causes and the use of the sinks by large numbers of people. Therefore, the aim of the present study was to verify the antibacterial efficacy of the moth-eye film by using hospital hand-wash sinks as practical environments for testing.

## Materials and Methods

This study was approved by the ethics committees of our university.

### Materials

Three types of sample films were used: a polyethylene terephthalate (PET) film as control; Moth-eye film, consisting of PET film coated with a nanoscale cone-shaped pillar surface made of hydrophilic resin; and Flat film, consisting of PET film coated with a flat surface made of hydrophilic resin. The Moth-eye coating was fabricated by the nanoimprint method using an ultra-violet curable resin and a Moth-eye stamper that possessed arrays of nanoscale cone-shaped holes [10]. The single unit structure of Moth-eye was a cone-shaped protrusion that was an inverted shape of the unit structure on the stamper that consisted of a hole that was approximately 200 nm in depth and diameter. To generate the Flat film, the Moth-eye stamper was replaced with a flat glass stamper surface.

### Laminating sample films

Sample films, each consisting of a 4-cm-square section of the respective material, were pasted onto the surface of the respective sink on a vertical surface inside of the sink and on a horizontal surface at the fringe of the sink. The arrangement of sample species were changed for each test in order to prevent the number of collecting colonized from being influenced from the pasting place. To adjust the test start time, after laminating, each of the sample film surfaces and each sinks were disinfected by wiping three times with an ethanol-impregnated paper cloth, and that time was taken as the start time of each tests.

### Collecting bacteria

Culture media stampers were used to collect bacteria. Each sample film was stamped by one stamper, using stampers smaller than the size of the sample film. The contact time of the stamper on the film was approximately three seconds. Three types of stampers, plate count agar (PCA) made of standard agar medium, mannitol salt with egg yolk agar (MSEY), and *Escherichia coli* (ES) Colimark agar (ESCM) were used (Eiken Chemical Co. Ltd., Japan).

### Testing locations

A total of 6 hand-wash sinks in three bathrooms (two sinks in each bathroom) were chosen at the university hospital. Numbers of sample films were 108 (48 Moth-eye, 48 PET, 12 Flat). The sinks were used constantly from 10:00 to 16:00, which correspond to the examination time. The time points were designed to collect bacteria at 1 hour and 6 hours after start time on the respective day. After the collection, the culture media stampers were incubated and numbers of colonies were counted.

An additional test was conducted at the company cafeteria, since the numbers of samples were too low to discuss statistically significant difference. A total of 4 hand-wash sinks was chosen at the company cafeteria. Numbers of sample films were 92 (46 Moth-eye, 46 PET). The sinks were used intensively during lunch break, from 11:30 to 13:00. The start time was 11:00, and the time points were designed to collect bacteria at 2 hours (13:00) after start time on the respective day. Following collection, the culture media stampers were incubated and numbers of colonies were counted.

To ensure fairness of the examination, the sampler who collected bacteria with stampers and the measurer who counted number of colonies were assigned separately. Therefore, the measurer counted without knowing the detail information on the sample films.

When counting, for all cases where the colonies were too numerous to count accurately, the number was defined as too-numerous-to-count (TNTC) and set at 400 for averaging purposes. Colonies that covered a relatively large area in the agar were omitted from the counts, and they were excluded from the verification.

### Statistical data processing

All data are reported as the mean and standard deviation. A two-tailed Mann-Whitney test was used to compare the effects of the individual films. A p-value less than 0.05 was considered statistically significant.

### Result and Discussion

The mean counts of live bacteria collected by PCA stamper from the three films are shown in Fig. 1. For both the 1-and 6-hour time points, the mean counts of common bacteria collected from Moth-eye and Flat were significantly lower than those from PET (p<0.05). In comparison between Moth-eye and Flat, those from Moth eye also were significantly lower than those from Flat at the 1-hour time point (p<0.05) but not at the 6-hour time point. These results mean that Moth-eye has two interesting properties in practical setting; immediate effectivity and durability of antibacterial effect.

**Fig. 1.**
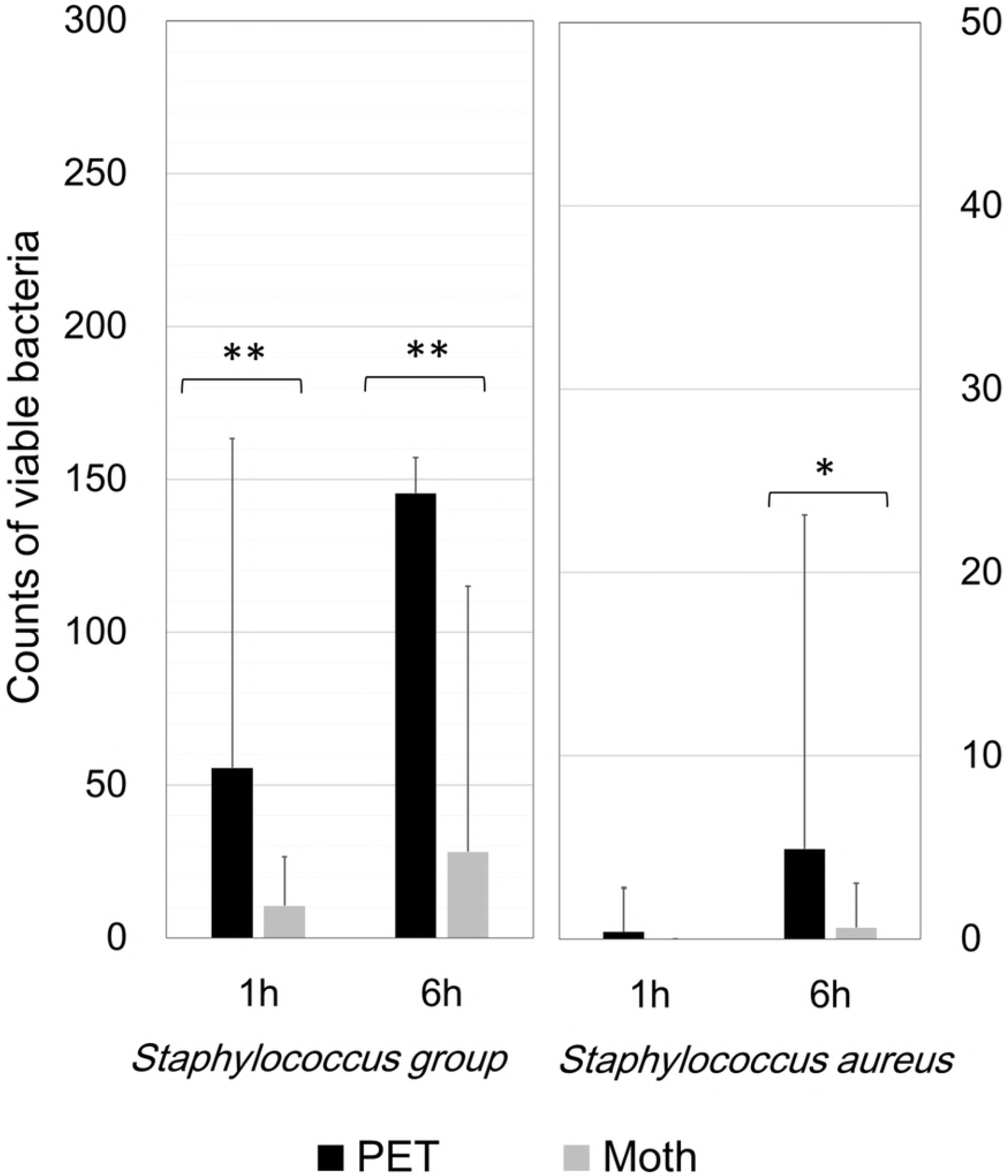
Counts of common bacteria on PCA stamper at the university hospital bath rooms. Average numbers of viable bacteria collected from the Moth-eye, Flat, and PET films after 1 hour and 6 hours. ** p<0.05, Mann-Whitney test. n=24.

Firstly, immediaye effectivity of Moth-eye was considered that nano-structure with hydrophilic resin enhanced the antibacterial effect, which was reported in our previous *in vitro* study [9]. Furthermore, the super-hydrophilic property of Moth-eye leads to a quick drying of water droplets that adhere to the moth-eye surface. This drying effect was expected to make it difficult for bacteria to grow on this surface.

Secondly, it was confirmed that the antibacterial effect of Moth and Flat with the surface material, hydrophilic resin, were kept longer than that of PET. Alcohol, well known as a disinfective chemical agent, has an immediate effect, but the effect is rapidly lost via evaporation of this reagent. It has been reported that the recovery of viruses and bacteria from stainless steel surfaces at 1 h after cleaning with ethanol ranged from 24 to 76% [11]. Most solid disinfective chemical agents exhibit their effects by diffusion out into water; antibacterial efficacy of these agents is lost once all of the agent has eluted. Efforts to provide longer durability have included adsorbing silver ions onto zeolite [12] and maintaining high concentrations of silver ions by using a super-hydrophilic binder [13]. At the same time, chemical agents are known to have negative impacts or risks, notably via the killing of useful microbial organisms and via the selection of drug-resistant bacteria.

The mean counts of live bacteria collected by MSEY stamper from Moth-eye and PET are shown in Fig. 2 and those by ESCM are shown in Fig. 3. Regarding the *S.* group and *S. aureus*, the numbers of colonies collected from Moth-eye were significantly lower than those from PET (p<0.05).

**Fig. 2.**
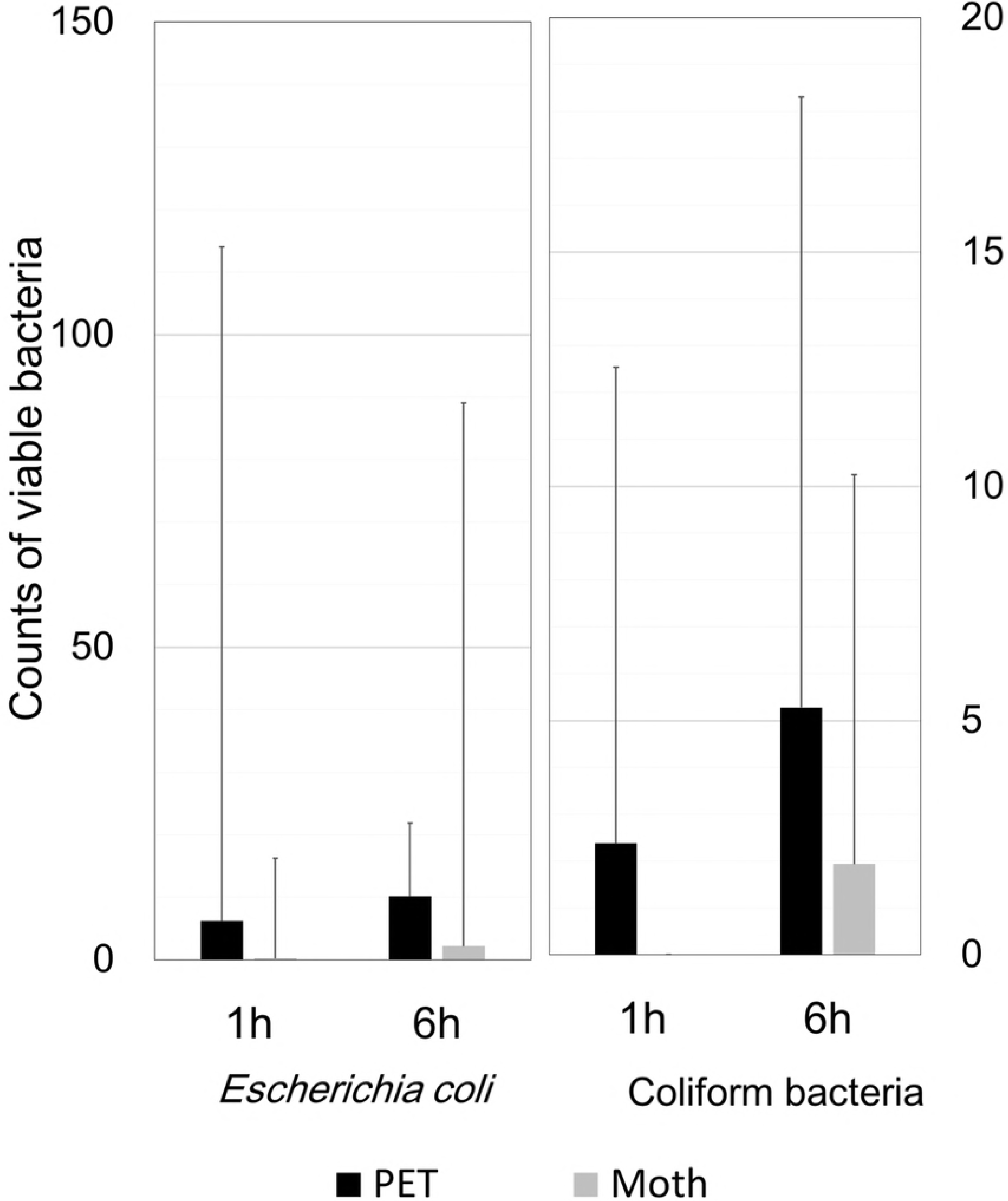
Counts of *Staphylococcus* group on MSEY stamper at the university hospital bath rooms. Average numbers of viable bacteria, which were members of the *Staphylococcus* group (a) or *Staphylococcus aureus* (b), collected from the Moth-eye and PET films after 1 hour and 6 hours. ** p<0.05, Mann-Whitney test. n=40.

**Fig. 3.**
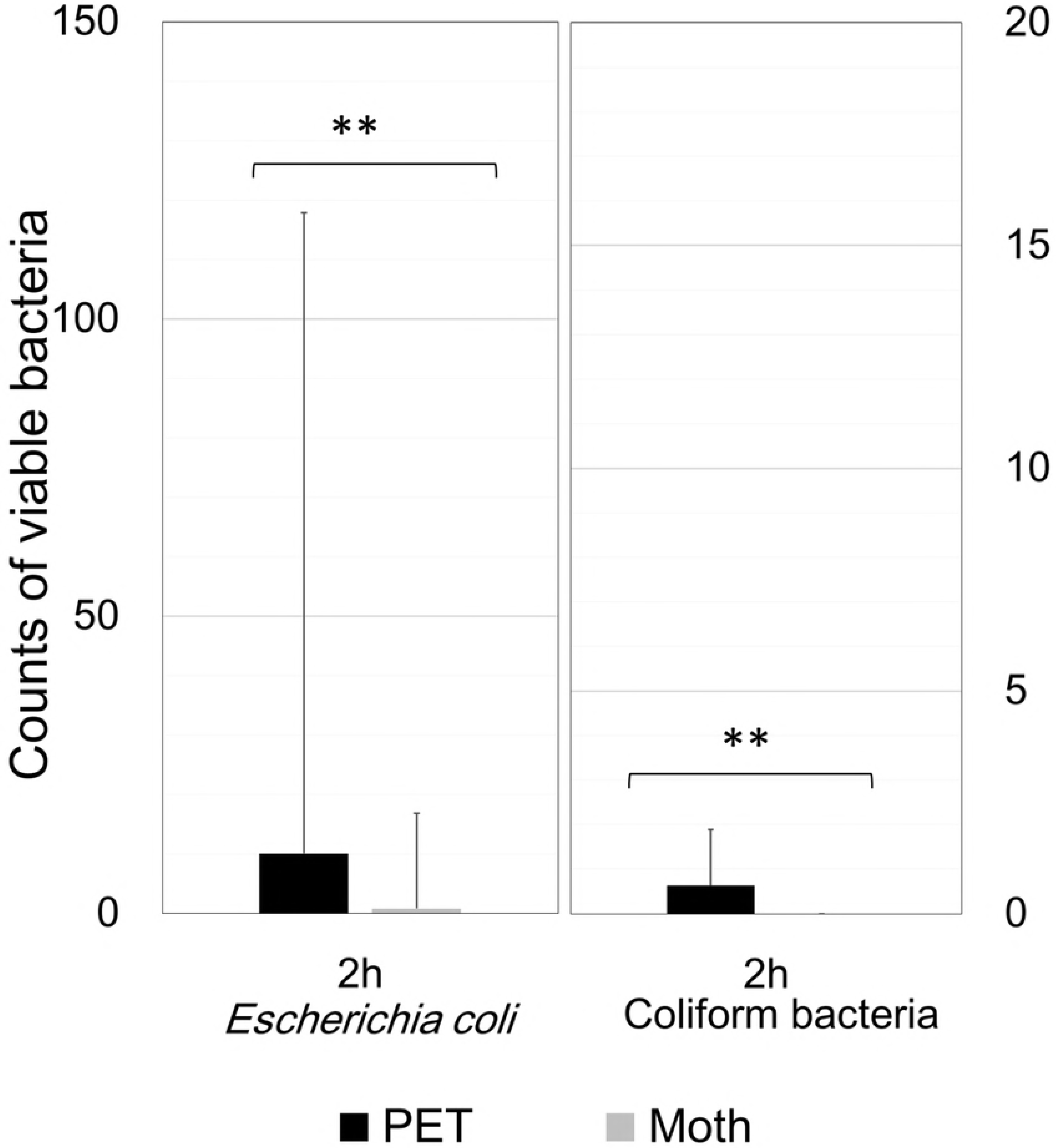
Counts of *Escherichia coli* and coliform bacteria on ESCM stamper at the university hospital bath rooms. Average numbers of viable bacteria, which were *Escherichia coli* (a) or coliform bacteria (b), collected from the Moth-eye and PET films after 1 hour and 6 hours. The numbers of bacteria did not significantly differ between the two films at either time point. n=32.

However, for both *E. coli* and coliform bacteria, the average counts from Moth-eye were nominally lower than those from PET, but the differences fell short of significance, possibly due to high variation. To collect more samples, we carried out the study at the company cafeteria and the results are shown in Fig. 4. As for both *E.coli* and coliform bacteria, the mean counts collected from Moth-eye were significantly lower than those from PET (p<0.05). These results suggested that Moth-eye might be useful for inhibiting the growth of bacteria such as *S. aureus* and *E.coli*, which are species associated with highly pathogenic hospital infection.

**Fig. 4.**
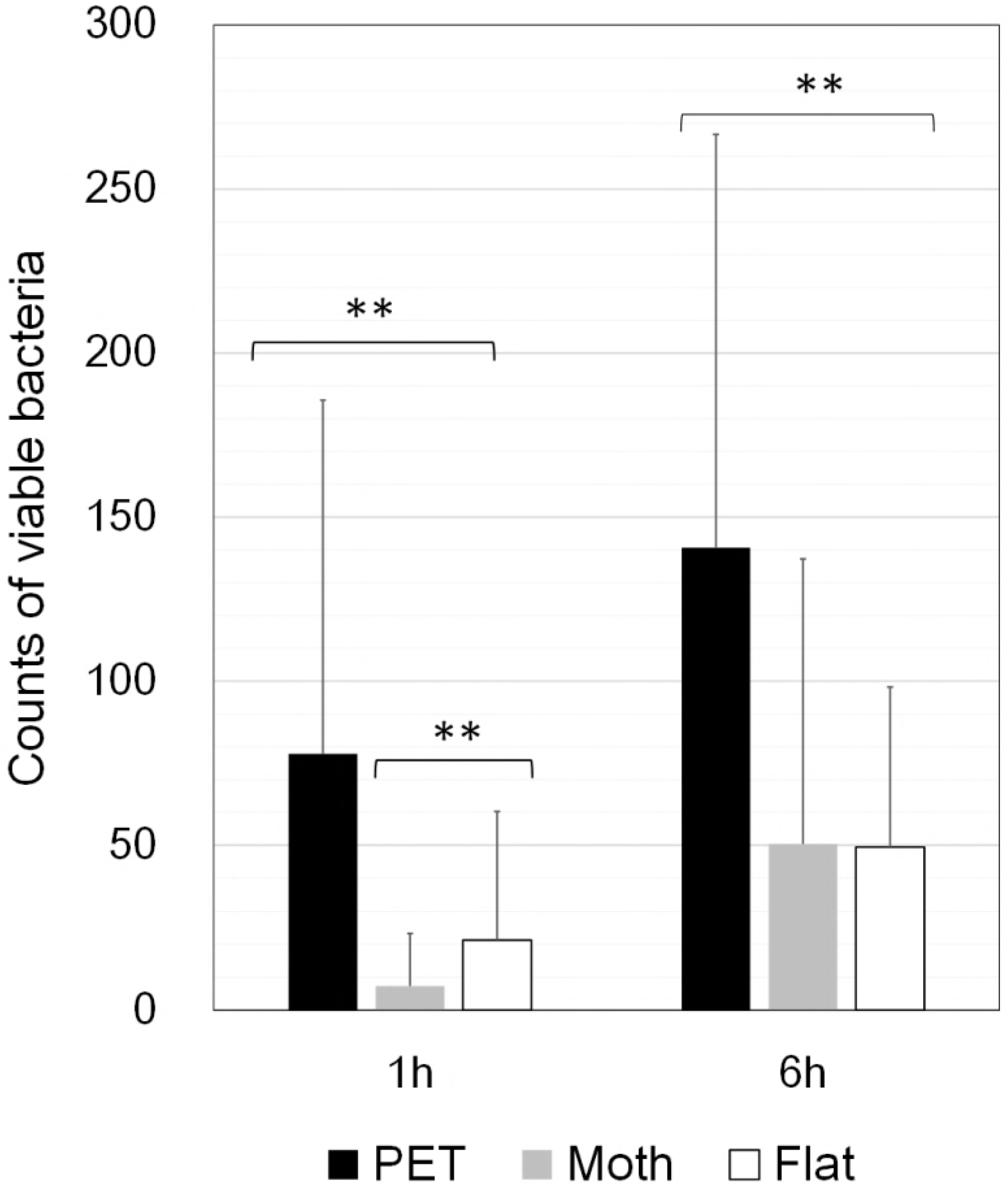
Counts of *Escherichia coli* and coliform bacteria on ESCM stamper at the company cafeteria. Average numbers of viable bacteria, which were *Escherichia coli* (a) or coliform bacteria (b), collected from the Moth-eye and PET films after two hours. ** p<0.05, Mann-Whitney test. n=46.

This study found an antibacterial effect for Moth-eye film deployed in public places, an effect that presumably included physical (nano-structure) and chemical (hydrophilic resin) properties. The safety of hydrophilic resin-coated PET had previously been confirmed to have no impact on rabbit skin by Animal Irritation test. The advantage of the Moth-eye film could include this material’s ability to bring about a decrease in contact infection risk by immediate antibacterial activity, along with longer-term suppression of bacterial growth. Moth-eye might be applicable to any equipment or surface in proximity to water, such as hand dryers, bathroom door knobs, water condensation trays, and so on. Further studies to investigate the durability of this film in various conditions will be needed to demonstrate if the Moth-eye film will be widely applicable for use in this context.

Based on the result of our previous study which showed difference of formation of colonies between on the nano structured surface and on flat surface, further studies in microscopic view point in practical setting will be investigated, whether to form biofilms or not for example.

